# Low Molecular Weight Hyaluronan Inhibits Lung Epithelial Ion Channels by Activating the Calcium-Sensing Receptor

**DOI:** 10.1101/2022.09.07.506946

**Authors:** Ahmed Lazrak, Weifeng Song, Zhihong Yu, Shaoyan Zhang, Anoma Nellore, Charles W. Hoopes, Bradford A. Woodworth, Sadis Matalon

## Abstract

Herein, we tested the hypothesis low molecular weight hyaluronan (LMW-HA) inhibits lung epithelial ion transport in-vivo, ex-vivo, and in-vitro by activating the calcium-sensing receptor (CaSR). Intranasal instillation of LMW-HA (150μg/ml) to C57BL/6 mice inhibited their alveolar fluid clearance (AFC) by 75%, increased the epithelial lining fluid (ELF) thickness threefold, and lung wet/dry (W/D) ratio by 20% 24hrs later. Incubation of lung slices from mouse and human lungs with 150μg/ml LMW-HA decreased the open probability (P_o_) of ENaC in ATII cell by more than 50% in 4hrs, inhibited amiloride sensitive short circuit current (SCC) 4hrs post exposure, and Cl^−^ current through CFTR by more than 70%, and Na,K-ATPase current by 66% at 24hrs. In all cases the inhibitory effect of LMW-HA on lung epithelial ion transport in vivo, ex vivo, and in vitro preparations were reversed by the administration of 1μM of NPS2143, a CaSR inhibitor, or 150μg/ml HMW-HA. In HEK-293 cells co-transfected with CaSR and the calcium sensitive Cl^−^ channel TMEM16-A, LMW-HA activated an inward Cl^−^ current. These data are the first demonstration of the inhibitory effects of LMW-HA on lung epithelial ion and water transport, and are due to the activation of CaSR and its downstream signaling cascades.

## Introduction

One of the hallmarks of lung injury is the breakdown of the cell matrix hyaluronic acid (HA)[1] to low molecular weight fragments [2] that are inflammatory [3]. Their accumulation in the distal lung is also associated with fluid accumulation and lung edema because of their ability to attract and retain water [4, 5]. HA interacts through hydrogen bonds, van der Waals, and electrostatic forces with hyaladherins, and with membrane receptors to modulate development, morphogenesis, cell migration, apoptosis, cell survival, inflammation, and tumorigenesis [6-11]. Reactive oxygen species (ROS) and hyaluronidases were found to dissociate HA into smaller fragments of low-molecular-weight (LMW-HA)[12-15]. LMW-HA binds to its membrane receptors, CD44, TLR4, and RHAMM and triggers intracellular signaling cascades that affect multiple cellular functions. HA interaction with ion channels is still understudied. In two studies, it was found to inhibit the pain sensitive channel TRPV1 [16], and reduce the open probability of a stretch-activated ion channel heterologously expressed in Xenopus oocytes[17]. LMW-HA was found to induce lung hyper-reactivity by increasing Ca_i_^2+^ in airways smooth muscles through the activation of G-protein coupled CaSR [18-20].

CaSR, as its name suggests, is a membrane receptor that senses and regulates the extracellular calcium concentration by controlling the release of the parathyroid hormone (PTH) [21-23]. Hence, CaSR is pharmacologically targeted for the treatment of mineral metabolism disorders of hyperparathyroidism and hypocalcemia using activators or calcimimetics, and inhibitors or calcilytics, respectively. In addition to its expression in organs involved in calcium homeostasis, parathyroids, kidneys, intestines, and bone, CaSR is also found in tissues with no role in calcium metabolism such as airway smooth muscle cells, blood vessels, breast, and placenta [20, 24-28]. CaSR enhanced expression and function were linked to pathological conditions such as inflammation [29-31], airways hyper-responsiveness [20, 32], cancer [33-35] and Asthma [25]. The primary ligand of CaSR is Ca^2+^, however, in the non-calciotropic tissues its function and expression have been found to be modulated by numerous other ligands including protons, polyvalent cations, amino acids, polyamines, polypeptides, antibiotics, and even anions [36-42]. Its ability to respond to such a diverse array of stimuli would make it relevant to a number of respiratory system pathophysiological conditions.

Yet, there is currently no evidence of CaSR involvement in ARDS, lung edema, or vectorial ion transport across lung epithelia. Consequently, we hypothesized that if CaSR was to be found in lung epithelia, it would sense and respond not only to increased [Ca^2+^]_o._ but also to polycations, polyamines, and anions such as LMW-HA, whose production is markedly increased during lung injury. To test our hypothesis, we used LMW-HA to mimic lung injury and study the role of CaSR on AFC, W/D ratio, the thickness of the epithelial lining fluid, the function ENaC in AT2 cells in precision cut lung slices from both mice and human lungs, and the function and the expression of ENaC, CFTR and the Na,K-ATPase in confluent and differentiated MTEC monolayers. To the best of our knowledge, this is the first study examining the role of CaSR in lung epithelial ion and water transport deregulation and lung edema formation.

## Methods

### Reagents and supplies

HMW-HA (Yabro) a gift from IBSA Institut Biochimique, Lugano, Switzerland; LMW-HA was generated by sonication of Yabro.

Antibodies: αENaC (Invitrogen PA1-920A), βENaC (proteintech 14134-1-AP), γENaC (Invitrogen PA577797), CFTR (abcam ab2478), Na,K-ATPase-α (abcam ab76020), Na,K-ATPase-β1 (santa cruz sc-21713), CaSR (Sigma-Aldrich C0493); actin (Sigma-Aldrich A5228).

Media and compounds used for MTEC isolation and culture: Ham’s F12 media (Corning, MT10080CV); DMEM/F-12 medium (Corning, 11-320-033); Keratinocyte SFM (Gibco, 17005042); 100 X Antibiotic-Antimycotic (Pen/Strep/Fungiezone) Solution (Cytiva HyClone, SV3007901); Pronase (Roche, 10165921001), DNAse I (Sigma Aldrich, 10104159001); Murine EGF (Sigma, E5160); Bovine Pituitary Extract (Gibco, 13028014); Isoproterenol (Sigma, I-6504); Y-27632 (Selleck Chemical LLC, 129830-38-2); DAPT (Sigma, D5942); DMEM/F12 (Gibco, 1133032); Fetal Calf Serum (HyClone, SH30071.03); 5% ITS-G (Gibco, 41400045); Cholera Toxin (Sigma, C8052); Retinoic Acid (Tocris Bioscience, 069550); BSA (Gibco, 15260037); Transwell™ Multiple Well Plate with Permeable Polyester Membrane Inserts (Corning, 07-200-154).

### Animals

C57BL/6 8 to 12 weeks old mice (20–25 g body weight), male or female, were purchased from Charles River Laboratories (Wilmington, MA). All experimental procedures involving animals were approved by the University of Alabama at Birmingham Institutional Animal Care and Use Committee (IACUC) Animal Project Number (APN): IACUC-22268.

### Human lungs

Human lungs that are unsuitable for organ transplantation were obtained from brain dead organ donors, after family had provided permission for organ retrieval for research purposes by the Legacy of Hope Organ Donor Center and enrolled into the UAB Non-Human Subjects Research protocol IRB-300004866. Human lungs were cannulated at the pulmonary artery and vein for 500 cc normal saline flush to remove all perfusate. The left upper lobe was resected for use in these studies.

### Nasal Potential difference (NPD) measurement

Adult C57BL/6 mice (8–10 weeks old; 20–25 g body weight), male or female, were exposed to chlorine (Cl_2_) gas (400ppm for 30 min) in cylindrical glass chambers as previously reported [20]. Control mice were exposed to air in the same experimental setup as Cl_2_. At the end of the exposure, the mice were returned to their cages and had access to food and water ad libitum. NPD measurements were performed 24hrs post exposure to Cl_2_ following the protocol published by Knowles et al.[43]. Mice were anesthetized by intraperitoneal injection of 100 mg/kg ketamine and 0.5 mg/kg dexmedetomidine. The recording cannula filled with Ringer’s solution was inserted in one of the mouse nostrils. The reference electrode (a needle filled with Ringer’s) was placed in the subcutaneous space of the hind leg. For NPD recording the nasal epithelium in close contact with the recoding cannula was perfused at rate of 200 μl/hr to allow a stable connection between the nasal epithelium and the recording amplifier (BMA-200 AC/DC Bioamplifier, CWE, Boston, MA). The recording was visualized on a monitor connected to a computer running Axopatch 10.6 (molecular devices, San Jose, CA) and interfaced with the amplifier using Digidata mini (molecular devices). When the recording cannula was inserted in the mouse nostril the sodium-driven NPD, a spontaneous downward deflection, of about -7 to -10 mV was recorded; it is indicative of the active sodium absorption through ENaC and inhibited by the addition of 200μM amiloride to the perfusate. After the complete inhibition of sodium driven NPD, the introduction of 20μM forskolin activated the chloride driven NPD through CFTR, −12 to −15 mV downward deflection in the recording trace, inhibited with 50μM GlyH-101.

### Lung wet/dry ratio (W/D)

To measure total lung water, lung weight is measured immediately after excision (wet weight), then after lung tissue is dried at 60°C for 48hrs and reweighted again. The wet/dry ratio, W/D, is calculated by dividing the wet weight of the lung by its dry weight[44]. For the experiment, control mice were instilled with 50μl saline, to test the effect of LMW-HA on lung W/D 50μl of 150μg/ml LMW-HA was instilled, then one hour later half of the mice instilled with LMW-HA were re-instilled with 50μl of 1μM NPS2143, to test the role CaSR in the retention of fluid in the lung following LMW-HA instillation. Twenty-four hours later animals were sacrificed and their lungs were surgically removed and weighted. To measure W/D of each group, lungs were placed in an oven at 60ºC degrees for 48 hours, then weighted again and the ratio was calculated.

### Alveolar fluid clearance (AFC) measurement

Twenty-four hours post 50μl of 10μg/ml LMW-HA instillation to mice AFC was measured as follow: **1**. Mice were anesthetized with an intraperitoneal injection of Ketamine 80-100mg/kg and xylazine 5-10mg/kg, depending on mouse weight. For reproducibility mice were paralyzed with an intraperitoneal injection of pancuronium bromide (0.04mg). **2**. The trachea was exposed and cannulated with an 18-gauge intravenous catheter trimmed to 0.5 inch long. **3**. Mice were positioned in the left decubitus position, and 0.5ml of 5% BSA was instilled over 30sec and infused with 0.1ml air in the catheter to clear the dead space and position the fluid in the alveolar spaces. **4**. Mice were ventilated for 30min to allow fluid clearance from the alveolar spaces. At the end of this period, the remaining alveolar fluid was collected using a 1ml syringe. **5**. AFC expressed as a percentage of total instilled volume, was calculated using the following formula: AFC = (1-Ci/C30)/0.95, where Ci and C30 are BSA concentrations at time zero and 30 min, respectively, measured using the bicinchoninic acid (BCA) protein assay[45].

### ELF thickness measurement

ELF thickness in mice instilled with 50μl of 150μg/ml LMW-HA was measured using micro-optical coherence tomography (μOCT). μOCT is a high-speed and high-resolution interferometry-based reflectance imaging modality, it was used to interrogate functional microanatomic parameters of excised trachea. μOCT resolution is less than one micron, which is sufficient to fully resolve ELF depth without the use of exogenous dyes or particles[46-48]. Trachea images were acquired at a rate of 40 frames per second at a resolution of 512 lines per frame. The imaging optics axis was placed within 10 degrees of normal to the tissue plane to reduce inaccuracies in geometric measurements. Three regions of interest per each trachea were documented and averaged for a single value per trachea. ELF thickness was calculated by direct measurement of the visible depth within the image. To account for refractory properties of the liquid, layer thickness measurements was corrected for the refraction index of the liquid (n=1.33). ELF depth was calculated from <u>pixels</u> conversion to <u>micro-meters</u> using a ratio of 0.82. Statistics were performed for each individual trachea replicates. All images were analyzed using ImageJ version 1.50i (National Institutes of Health, Bethesda, MD).

### Recording ENaC activity in human and mouse ATII cells in-situ

Lung slices were prepared from mice instilled with 50μl saline (control) or 50μl of 150μg/ml LMW-HA. Twenty-four hours post instillations, mice were euthanized using a lethal dose of Ketamine and Xylazine, then their chests were cut open, the lungs were washed with Krebs-Ringer solution containing (in mM): 140 NaCl, 3 KCl, 2.5 CaCl_2_, 1 MgCl, 10 glucose, and 10 HEPES (pH ≈7.35–7.4), and subsequently filled with low temperature melting agar through an incision in the trachea to stiffen the tissue for slicing with a microtome (Precisionary Instruments. Greenville, NC). Lower lobe of the right lung was dissected and mounted on the microtome and sliced into 250μm thick slices. For human lung, the left upper lobe was dissected and filled with low temperature melting agar to stiffen the tissue before slicing. The slices from human and mouse lungs were incubated in DMEM culture media at 37°C in humidified atmosphere of 5% Co_2_. For the experiment, a slice was transferred to the recording chamber filled with a solution of the following composition (in mM): 130 K-gluconate, 2 MgCl_2_, 10 KCl, 10 glucose, and 10 HEPES (pH 7.4, KOH) and held in place with an anchor (Warner Instruments). The recording pipette resistance ranged from 5-6 MΩ when filled with a solution of the following ionic composition (in mM): 140 NaCl, 3 KCl, 2.5 CaCl_2_, 1 MgCl, 5 glucose, and 10 HEPES (pH = 7.4). For ENaC activity measurement, the recording chamber was mounted onto the stage of an upright Olympus microscope EX51WI (Olympus, Center Valley, PA). To assess ENaC activity in ATII cells by patch clamp technique[49]. ATII cells within the slices were visualized by the accumulation of LysoTracker green (1 μM, ThermoFisher). Data were recorded and stored onto the hard drive of a computer equipped with pClamp software (Molecular Devices, San Jose, CA) and interfaced to an Axon amplifier Axopatch 200B (molecular devices) with Digitata 1440A (Molecular devices). Data were analyzed using Clampfit (molecular devices). The experimental groups were compared using ordinary ANOVA, p<0.05 was considered significant.

### Mouse tracheal epithelial cells (MTEC) isolation and culture

Four Mice were euthanized by an overdose of Ketamine/Xylazine (25μl of 100mg/ml and 2.5μl of 100mg/ml, respectively), the tracheae were removed, cut from the thyroid cartilage to the bifurcation, stripped of attached tissues, cut open lengthwise, washed in sterile PBS at room temperature for 5 min, then transferred to Collection medium (1:1 mix of DMEM: Nutrient Mixture Ham’s F-12 medium, 1% penicillin–streptomycin). Culture technique was as described previously [50, 51]. Batches of four tracheas were incubated in dissociation medium (collection medium plus 1.4 mg/ml Pronase, and 0.1 mg/ml DNase) overnight at 4°C. Next morning, epithelial cells were dispersed by gentle agitation, pooled, washed, and incubated in culture medium at 37°C for 2 h in a Premaria culture dish (Becton Dickinson, USA) to remove the non-epithelial cells. Non-adherent cells were collected, washed, and seeded in a T75 flask and incubated at 37°C in 5% CO_2_ atmosphere in a humidified incubator until confluent. Cells were then dissociated and seeded on permeable support in growth medium (Collection media with 2% Ultroser-G serum substitute) until they form tight confluent monolayers of differentiated tracheal epithelial cells.

### Bioelectric properties of MTEC

Confluent and differentiated MTEC monolayers were studied in bilateral Krebs-Ringer solution of the following composition (in mM): 120 NaCl, 25 NaHCO_3_, 3.3 KH_2_PO_4_, 0.83 K_2_HPO_4_, 1.2 CaCl_2_, 1.2 MgCl_2_, 10 HEPES (Na^+^-free), and 10 glucose (basolateral compartment). Bath solutions were mixed continuous bubbling with 95% O_2_, 5% CO_2_ at 37°C (pH 7.4). MTEC monolayers were mounted in Ussing chambers connected to a current/voltage clamp amplifier (Physiological Instruments, San Diego, CA.), All measurements were performed under short-circuit current (SCC) conditions. Currents measurement allowed the assessment of each pathway function using specific activator and inhibitors. We used amiloride (10μM) to measure the current carried by Na^+^ through ENaC, after reaching a steady state, we applied 10μM Forskolin to activate CFTR. To confirm the activated current is through CFTR, we used 20μM CFTR-_inhib.-172_ to block CFTR channels. Na,K-ATPase maximal activity was assessed when cells were loaded with Na^+^ following the permeabilization of MTEC apical membranes with 100μM amphotericin B. To confirm the current resulted from Na,K-ATPase activity, 200μM ouabain was added to the basolateral side of monolayers to inhibit the pump activity. SCC was assessed at 4- and 24-hrs post 50μl of 150μg/ml LMW-HA application to the apical membranes of confluent and differentiated monolayers. To reverse the inhibitory effects of LMW-HA on ion channels and the pump, we applied 50μl of 150μg/ml HMW-HA or 50μl of 1μm NPS2143, CaSR inhibitor, at 6hrs post monolayers exposure to 150μg/ml LMW-HA and measured SCC at 24hrs.

### Whole cell recording

Current-voltage (I/V) relationships of Cl^−^ current through TMEM-16 stably expressed in HEK-293 cells were recorded using the whole cell mode of the patch-clamp technique [49]. During the experiments, cells were perfused at a rate of 1 ml/min with an external Ringer’s solution of the following ionic composition (in mM): 145 CsCl, 2 MgCl_2_, 2 CaCl_2_, 5.5 glucose, 10 HEPES, pH 7.4 (1 N CsOH). The pipette resistance used for whole cell recording ranged from 2 to 3 MOhms when filled with the following solution (in mM): 135 CsCl, 10 KCl, 2 MgCl_2_, 0.1 EGTA, 5.5 glucose, 10 HEPES, pH 7.2 (1 N CsOH). All measurements were performed at room temperature. Data were recorded and stored onto the hard drive of a computer equipped with pClamp software (Molecular Devices, San Jose, CA) and interfaced to an Axon amplifier Axopatch 200B (molecular devices) with Digitata 1440A (Molecular devices). Data were analyzed using Clampfit (molecular devices).

### Total and membranes proteins

ENaC subunits, CFTR, Na,K-ATPase subunits, and CaSR total proteins in MTEC monolayers were assessed at 4- and 24-hrs post LMW-HA application to monolayers using Western Blotting technique. Commercially available antibodies to α-ENaC (H59, Santa Cruz), β-ENaC (sc-25354, Santa Cruz), and γ-ENaC (ab-3468, Abcam), CFTR protein (ab2478, Abcam) and Na,K-ATPase α-subunit (ab76020, Abcam) and β1-subunit (sc21713, Santa Cruz) were used to assess the expression of each ion channel. These measurements were repeated in MTEC monolayers incubated with LMW-HA and treated with HMW-HA (IBSA, Lugano, Switzerland) or NPS2143 ((Tocris, Minneapolis, MN) at 6hrs post LMW-HA application to reverse and inhibit, respectively, LMW-HA effects on ion channels and the pump.

### Statistical analysis

Data analysis and presentation were perform using GraphPad Prism 9.3 (GraphPad Software, San Diego, CA). Data were summarized as means ± SE. Statistical significance between groups was performed using ANOVA, followed by Tukey’s test for multigroup comparisons. *t*-test was performed when comparing two groups. *P* < 0.05 was considered significant.

## Results

### LMW-HA inhibits AFC and increase lung W/D ratio

Intratracheal instillation of 50μl of 150μg/ml LMW-HA, generated by sonication of HMW-HA (**Figure 1A)**, to mice, inhibited AFC by 65% at 24hrs (**Figure 1B**) compared to control values, and increased lung edema by a 55%, shown as an increase of lung W/D ratio (**Figure 1C**). In addition, 150μg/ml LMW-HA increased ELF thickness, measured using μOCT (**Figures 1D, E, and F**), threefold (**Figure1 G)**, from 14.9±2.1μm in control mice to 45.8±6.5μm in mice Instilled with LMW-HA. The damaging effects of LMW-HA on AFC, W/D ratio, and ELF thickness were reversed by the instillation of HMW-HA (50μl of 150 μg/ml) or NPS-2143, CaSR inhibitor, (50 μl of 1μM dilution) at 6hrs post LMW-HA instillation to mice.

**Figure 1:**
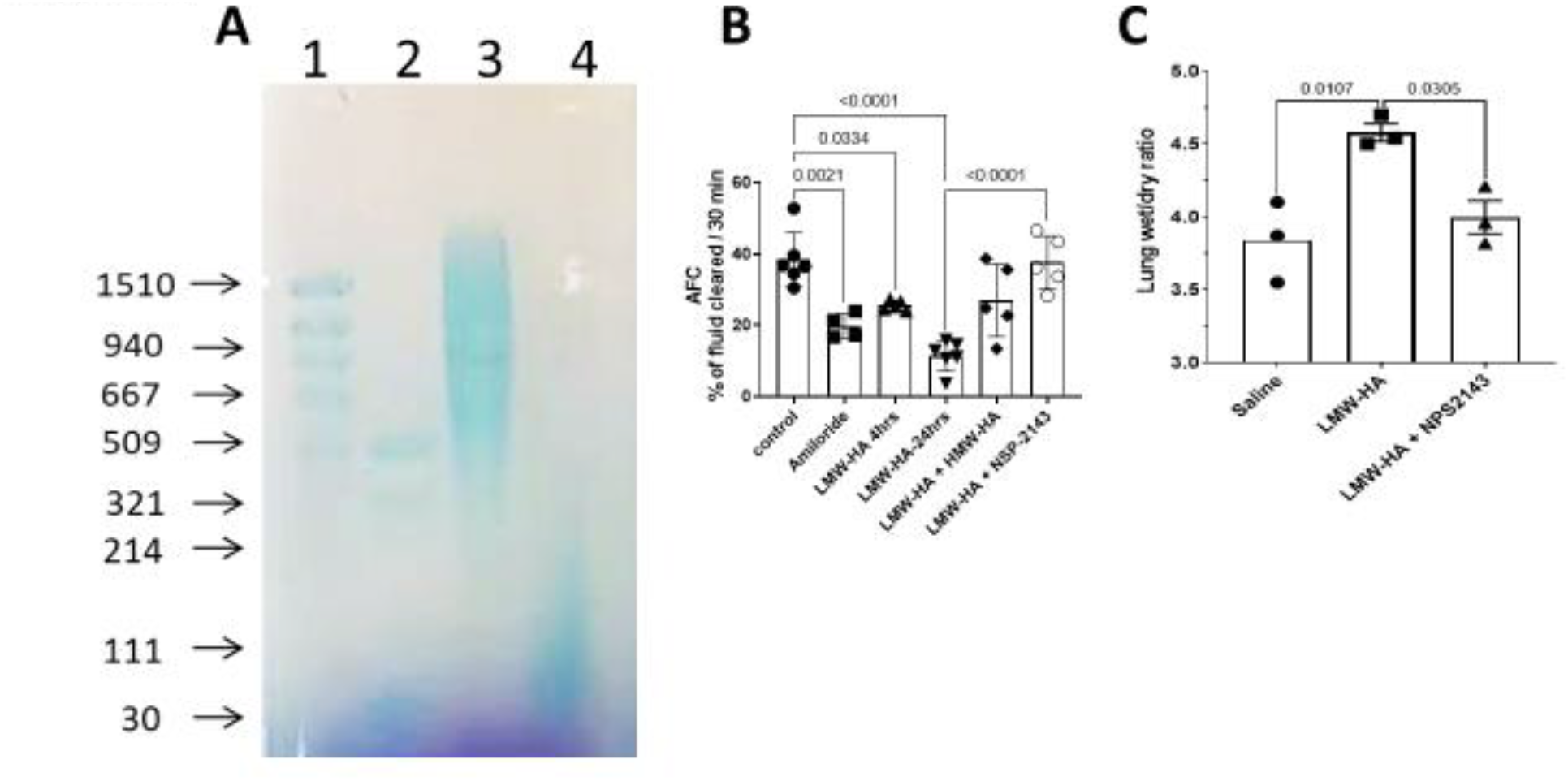
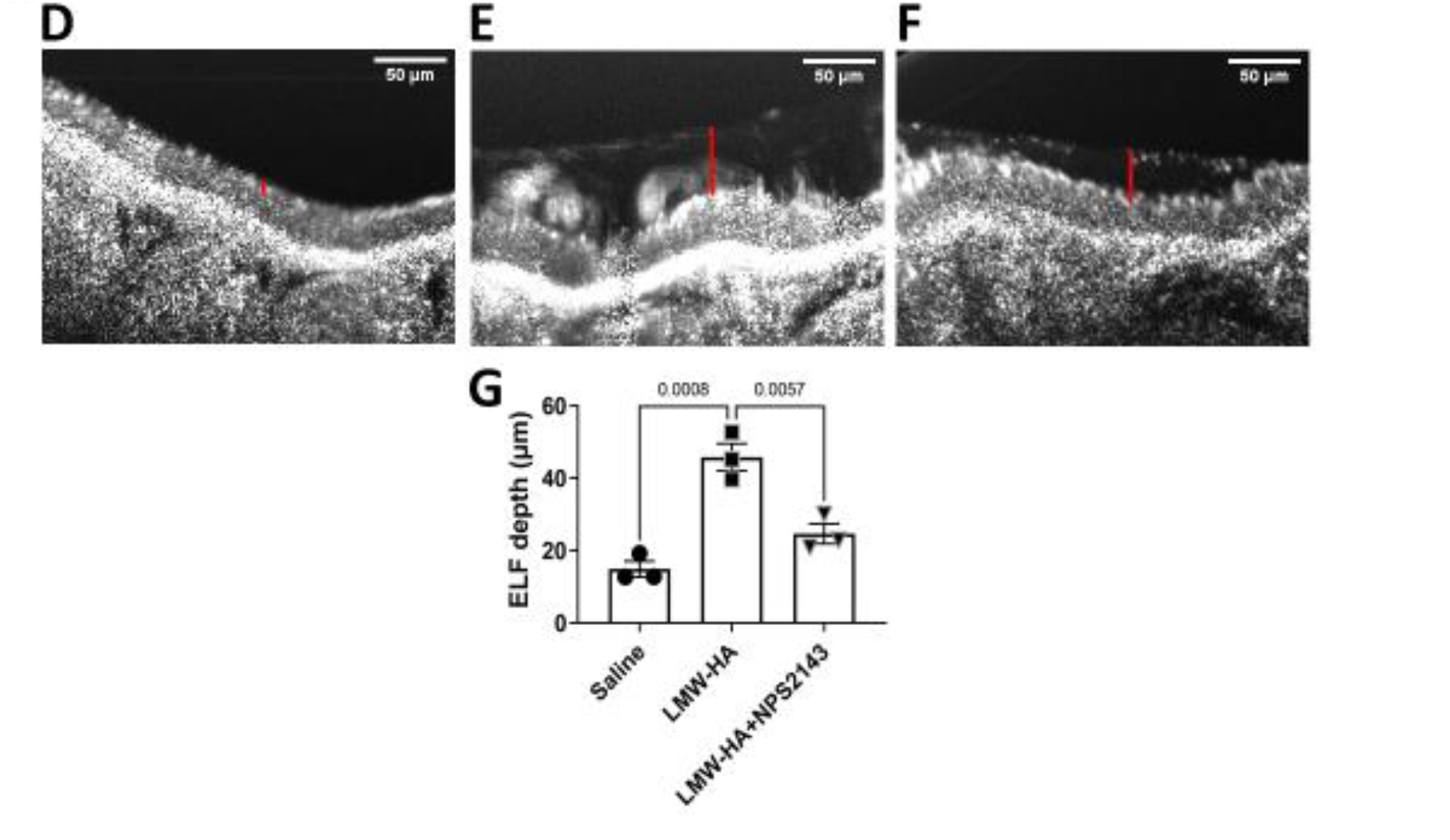
**A**. Agarose gel showing the breakdown of Yabro to lower molecular weight fragments (MW<500kD) by sonication for 1.5 min. The gel is stained with stain all. 1. HA-High ladder. 2. HA-Low ladder. 3. HMW-HA = Yabro. 4. LMW-HA. **B**. Instillation of 50μl of 150μg/ml LMW-HA to mice inhibited AFC starting at 4hrs and lasted at least 24hrs. Instillation of 50μl NPS2143 (1μM) at 6hrs post LMW-HA or 50 μl of HMW-HA (150μg/ml) reversed AFC to control values. **C**. Lung W/D ratio of 3 mice instilled with 50μl saline, 3 instilled with 50μl of LMW-HA at 150μg/ml, and 3 instilled with 50μl of NPS2143 at 1μM 6hrs post-LMW-HA. LMW-HA increased lung D/W 20%, while NPS2143 reduced it by 16%. **D, E** and **F**. Representative μOCT images of trachea from mice instilled with saline (D), 50μl of 150μg/ml HMW-HA (E), and 50μl of 1μM NPS2143 (F) at 6hrs post LMW-HA instillation. The red bar represents the thickness of the ELF. **G**. ELF increased threefold, from 14.9±2.1μm to 45.8±6.5μm, 24hrs post LMW-HA instillation. Inhibiting CaSR 6hrs post LMW-HA instillation with 1μM NPS2143 reduced ELF thickness by almost half, from 45.8±6.5μm to 24.8±4.8μm. Data are Means ± SE, n= 3-6. Significance was determined by 1-way ANOVA and post hoc Tukey test for multiple comparisons.

### LMW-HA inhibits ENaC activity in mouse ATII cells in-situ

Previously we have shown that more than 90% of AFC is secondary to the active transport of sodium ions across lung epithelial cells. Thus, the impairment of AFC and increase of ELF suggest that LMW-HA inhibited the activity of lung epithelial Na^+^ channels. Indeed, cell attached patches of ATII cells in precision cut lung slices revealed the presence of a highly selective sodium channel with a conductance of 4 pS (**Figures 2A-C**) and a non-selective cation channel with a conductance of 18 pS (**Figures 2D-F**), both amiloride sensitive, albeit with different EC50 values [52]. Instillation of 50μl of LMW-HA (150μg/ml) inhibited ENaC activity and reduced the open probability, P_o_, of both conductances as early as 4hrs of LMW-HA instillation (**Figures 2G&2H**). The inhibition persisted for at least 24hrs (Figs. 2G&H). Intranasal instillation of 50μl of HMW-HA (150μg/ml) or 50μl of 1 μM NPS2143 at 6hrs post LMW-HA administration reversed its effect on ENaC activity at 24hrs (Figs. 2G&H).

**Figure 2:**
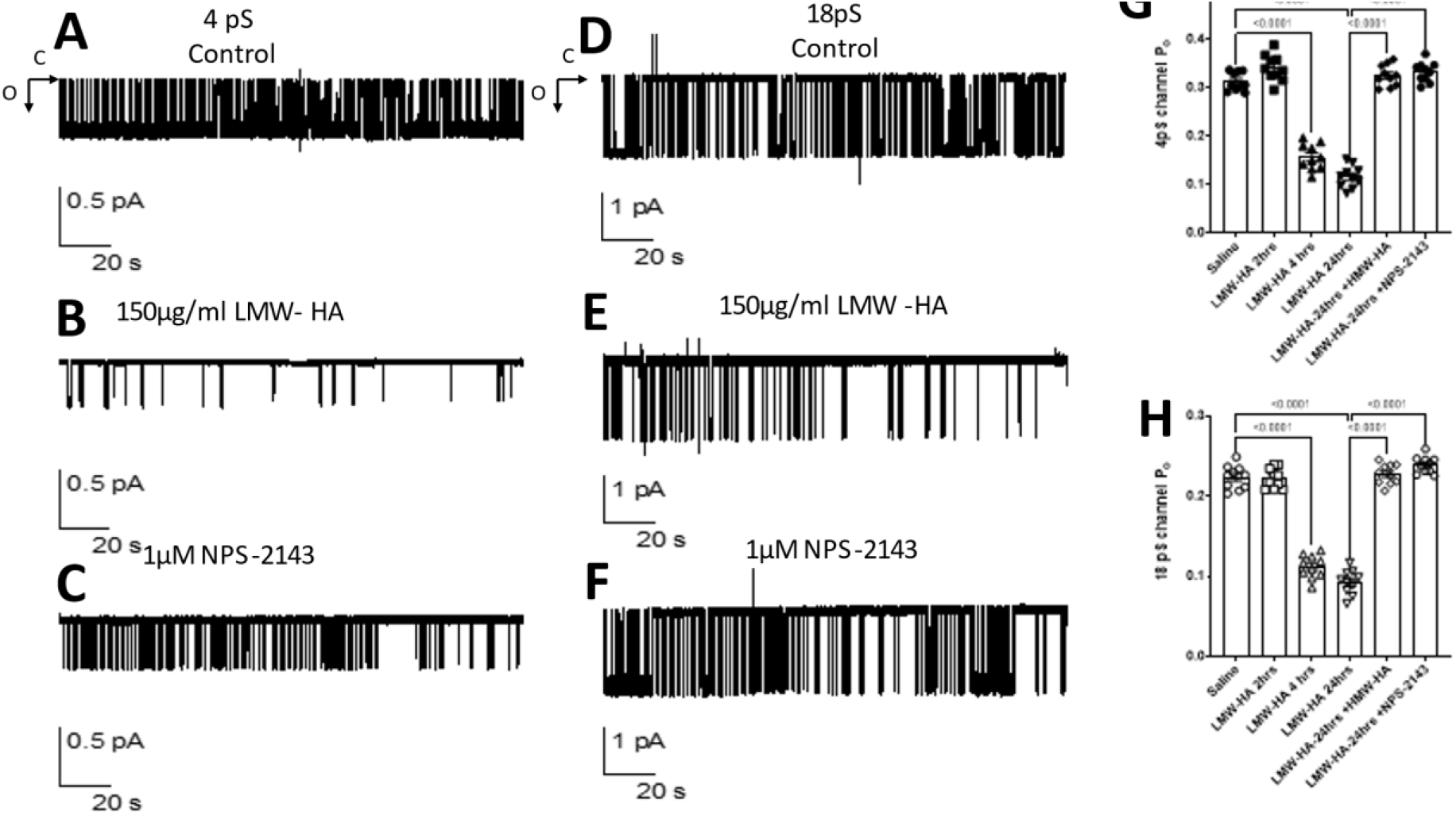
ENaC activity in mouse ATII cells in-situ showing two conductances 4pS and 18pS before and during exposure to LMW-HA (Panels 4A, 4B, 4D, and 4E), the arrows indicate the open (o) and closed (c) states of the channels. ENaC activity was restored when CaSR was inhibited with 1μM NPS-2143 (Panels 4C, and 4F). Panels 4G and 4H summarize the open probability of ENaC in AT2 cells in control, 2-, 4-, and 24hrs post LMW-HA instillation to mice. LMW-HA inhibitory effect on ENaC at 4hrs post LMW-HA instillation and was reversed only when CaSR was inhibited with 1μM NPS2143 instilled 6hrs post LMW-HA. Data are means ± SE, n = 10 slices prepared from three mice in each group. Significance was determined by 1-way ANOVA and post hoc Tukey test for multiple comparisons.

### LMW-HA inhibits short-circuit (SCC) in MTEC monolayers

Figure 3. illustrates representative SCC traces recorded across MTEC monolayers in primary culture mounted in Ussing chambers. **Panel A**, basal currents of both Na^+^, through ENaC, and Cl^−^ current, through CFTR, were inhibited by addition of 10 μM Amiloride or 20μM CFTR_inhib.-172_, respectively. Permeabilization of the apical membranes with amphotericin-B (100μM) activated an ouabain (200μM) sensitive current driven by the activity of Na,K-ATPase located at the basolateral membranes of monolayers. **Figure 3B**, illustrates the effect of 150μg/ml LMW-HA at 4hrs on SCC, LMW-HA inhibited amiloride sensitive current as early as 4hrs, without affecting Cl^−^ current through CFTR or Na,K-ATPase activity. At 24hrs post LMW-HA application all currents were inhibited, **Figure 3C & 3E**. Their function was restored to control levels when LMW-HA effect was reversed by the application of 50μl of 150μg/ml HMW-HA (**Figure 3F**), or 50μl of 1μM NPS-2143 **(Figure 3G)** at 6hrs post 50μl of 150μg/ml LMW-HA application to the apical membranes of MTEC monolayers without affecting monolayers resistance (data not shown).

### ENaC, CFTR, and Na,K-ATPase subunits protein expression

In the next series of experiments, we added 150μg/ml LMW-HA to the apical sides of MTEC monolayers and assessed the expression of ENaC subunits, CFTR, Na,K-ATPase subunits, and CaSR proteins at 4hrs and 24hrs later using specific antibodies (see Methods section). Characteristic blots are shown in **Figure 4A** and the quantification of the bands in **Figures 4-B-H**. LMW-HA addition to apical membranes decreased α-, β-, γ-ENaC, CFTR, Na,K-ATPase α and β_1_ subunits expression at both, 4hrs (Fig. 4A, lanes 3, 4) and 24hrs (Fig. 4A, lanes 5 and 6). Addition of either HMW-HA or NPS-2143 at 6hrs post LMW-HA application to MTEC monolayers restored ENaC subunits, CFTR and Na, K-ATPase protein expression to control levels (Fig 4A, lanes 7 and 8) and (Fig. 4A, lanes 9 and 10), respectively. In contrast, LMW-HA increased the expression of CaSR at 4hrs (Fig. 4A, lanes 3 and 4) and 24hrs (Fig 4A lanes 5 and 6), this effect was reversed by 150μg/ml HMW-HA (Fig. 4 Lanes 7 and 8) and 1μM NPS-2143 (Fig. 4 lanes 9 and 10) added at 6hrs post 150μg/ml LMW-HA apical application to MTEC monolayers.

**Figure 3:**
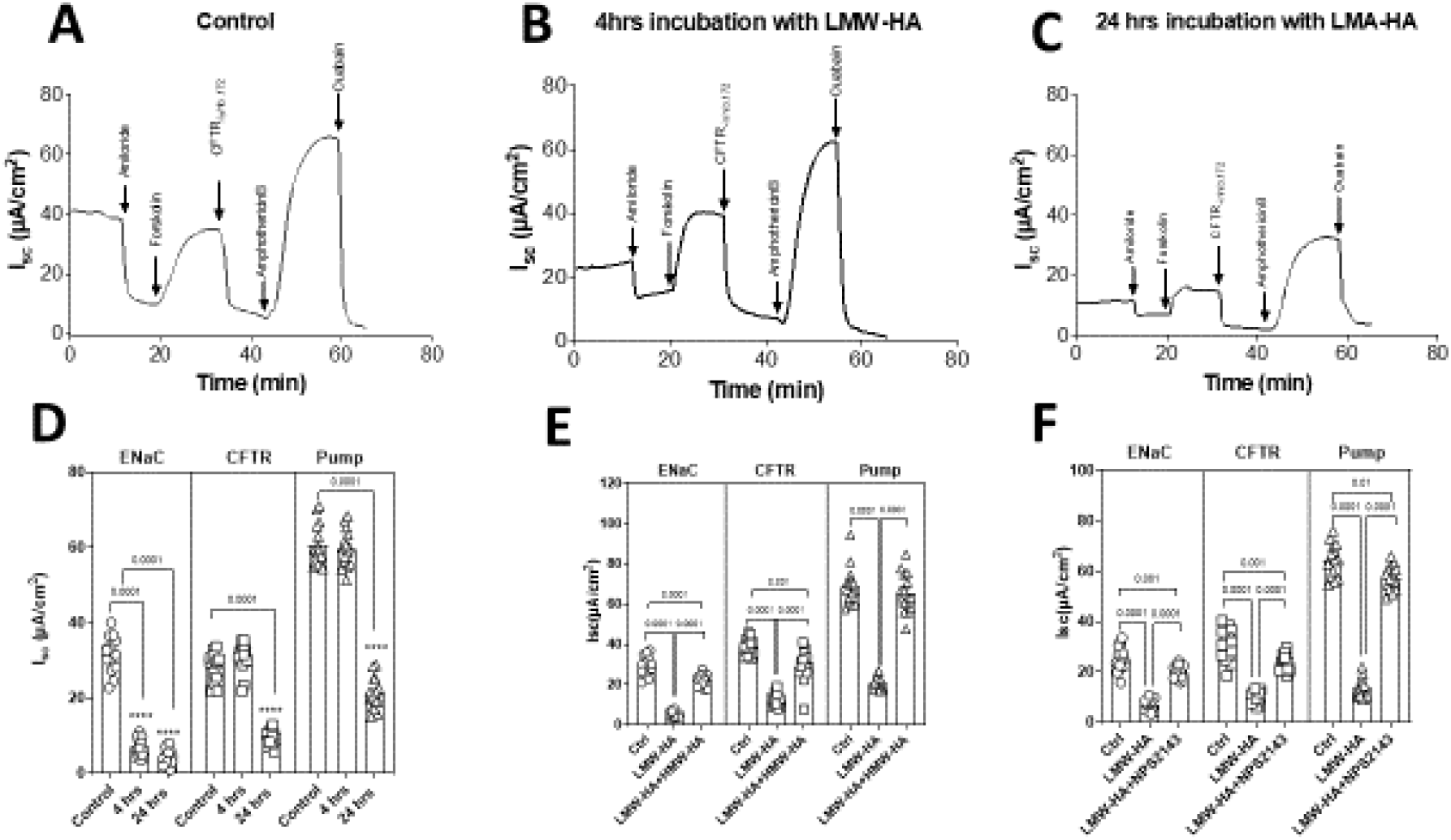
**A**. SCC recorded from MTEC monolayer. The basal current of 40μm/cm^2^ is mostly amiloride sensitive carried by Na^+^ ion entering the cell through ENaC. Forskolin (10uM) activated current and inhibited with CFTR-inhib-172 is a Cl^−^ current through CFTR. The Na,K-ATPase current visible when apical membranes are permeabilized with amphotericin-B and sensitive to ouabain. **B**. Apical application of 150μg/ml LMW-HA to MTEC monolayers inhibited the amiloride sensitive current as early as 4hrs without affecting the CFTR-_ihib.172_ or ouabain sensitive currents. **C**. 24hrs post incubation with 150μg/ml LMW-HA all currents were inhibited. **D**. Summary of SSC at 4hrs and 24hrs post MTEC monolayers incubation with 150μg/ml LMW-HA. ENaC is inhibited as early as 4hrs. At 24hrs, in addition to ENaC, CFTR and Na,K-ATPase, are also inhibited. **E**. LMW-HA inhibition of SCC at 24hrs was reversed by apical application of 150μg/ml HMW-HA at 6hrs post 150μg/ml LMW-HA application to MTE monolayers. **G**. CaSR inhibition with 1μm NPS2143 at 6hrs post 150μg/ml LMW-HA application to MTE monolayers reversed LMW-HA effect on SCC. Data are means ± SE, n= 13-15, p<0.001. Significance was determined by 1-way ANOVA and post hoc Tukey test for multiple comparisons.

**Figure 4:**
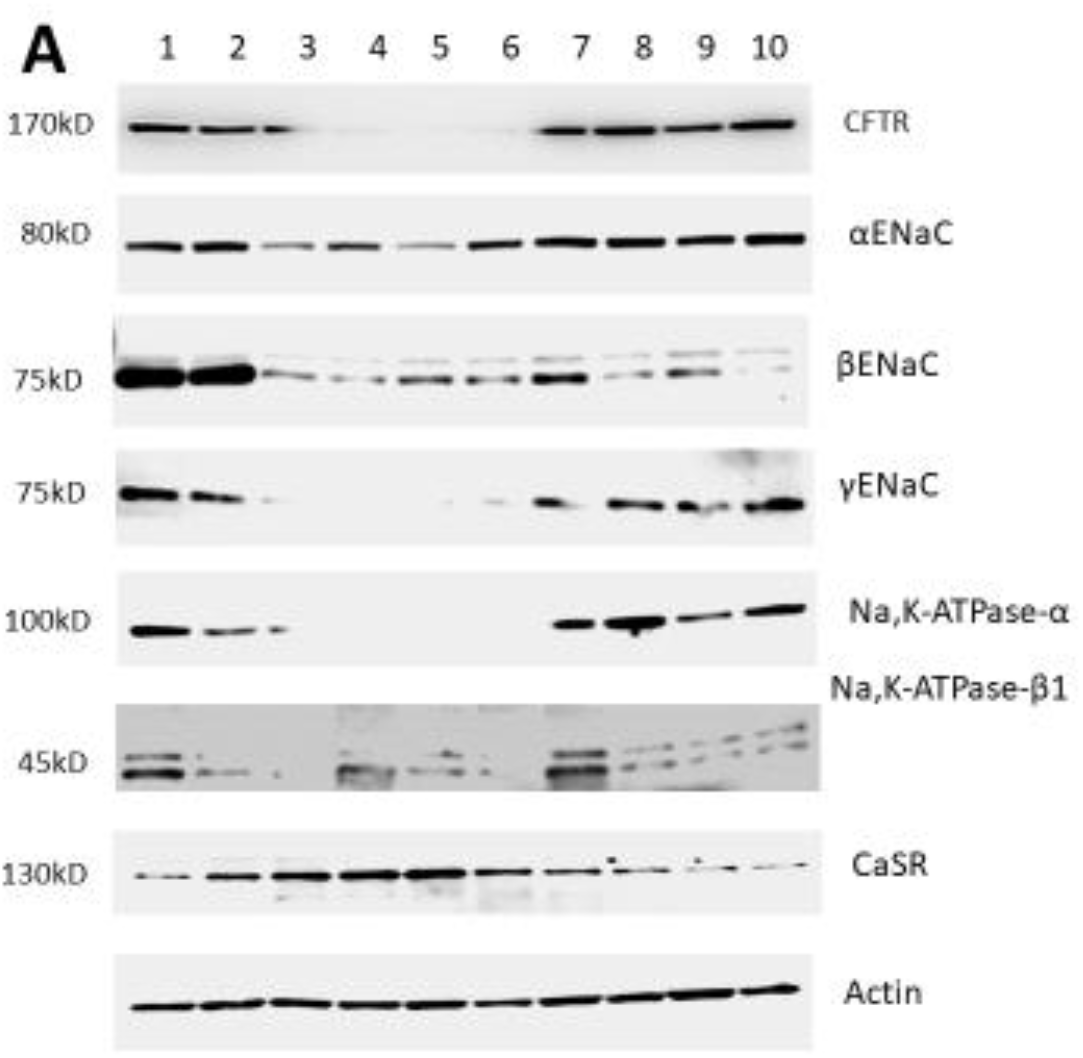
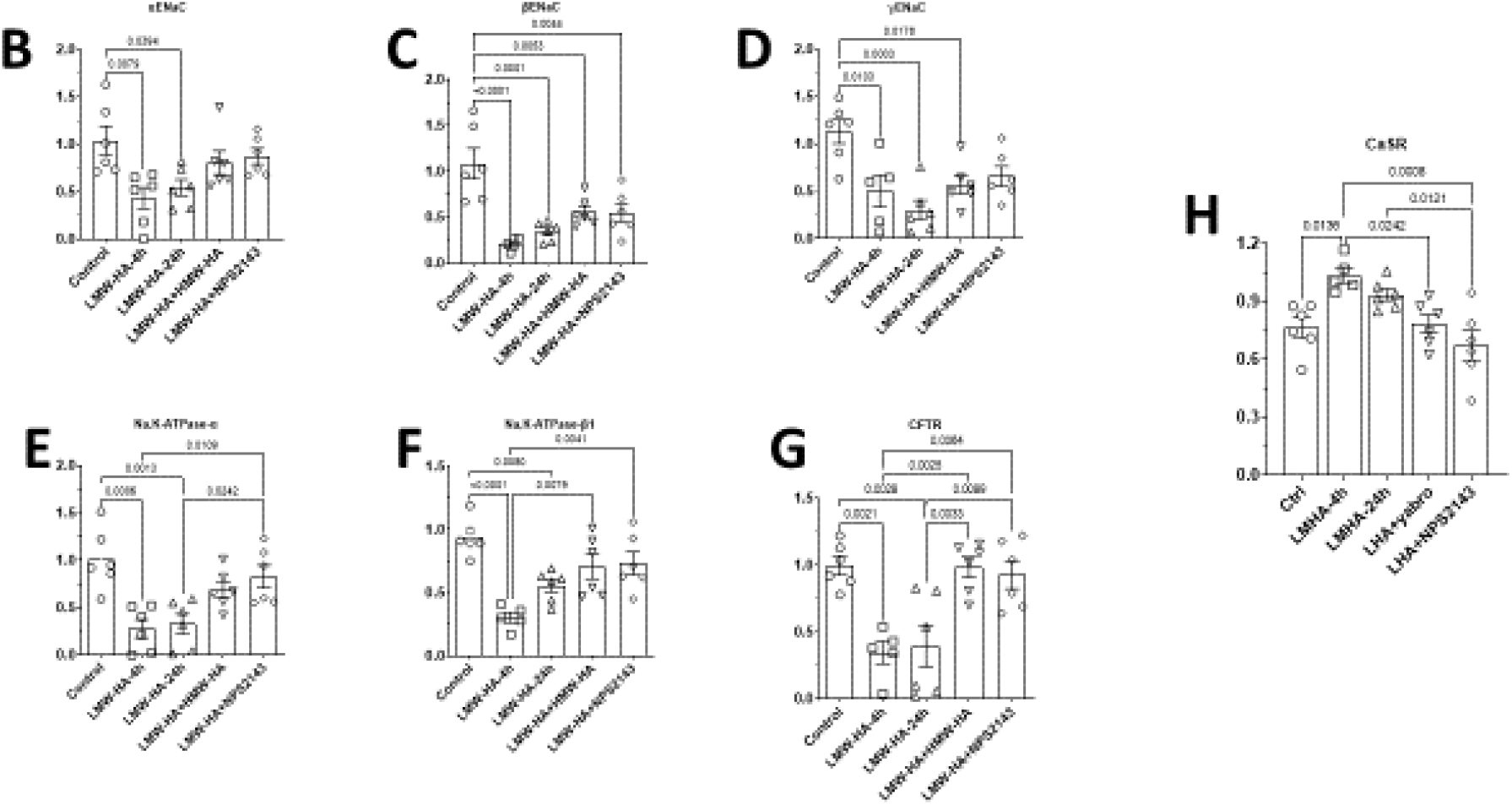
**A**. Total proteins for α-, β-, and γ-ENaC, CFTR, α-β-Na,K-ATPase subunits and CaSR. Control (lanes 1-2). 150μg/ml LMW-HA 4hrs (lanes 3-4). LMW-HA 24hrs (lanes 5-6), LMW-HA + 1μM NPS-2143 (lanes 7-8), and LMW-HA + HMW-HA (lanes 9-10). LMW-HA reduced ENaC subunits, CFTR, Na,K-ATPase subunits expression as early as 4hrs. However, it stimulated the expression of CaSR protein expression. 1μ M NPS-2143 and 150μg/ml HMW-HA restored ENaC subunits, CFTR, Na,K-ATPase subunits, and CaSR expression to their control levels. **B-H**. Quantification of western blot data in A for ENaC subunits, CFTR, Na,K-ATPase subunits and CaSR. Combined results of different 5 blots from 5 different MTEC cultures. Data are Means ± SE, n=5. Significance was determined by 1-way ANOVA and post hoc Tukey test for multiple comparisons.

### Inhibition of PLC, PKC and ERK abrogates LMW-HA effect on SCC across MTEC monolayers

To investigate downstream pathways involved in the inhibition of SCC by LMW-HA and CaSR, we pretreated MTEC monolayers with PLC, PKC or ERK inhibitors 4hrs prior to 150μg/ml LMW-HA application to monolayers and measured SCC at 4hrs and 24hrs. **Figure 5A**, shows the inhibitory effect of LMW-HA on amiloride sensitive current at 4hrs was reversed when either PLC or PKC were inhibited. ERK inhibition did not reverse ENaC function at 4hrs. However, at 24hrs the function of ENaC, CFTR, and Na,K-ATPase were restored when either PLC, PKC, or ERK were inhibited, **Figures 5B, 5C, and 5D**.

**Figure 5.**
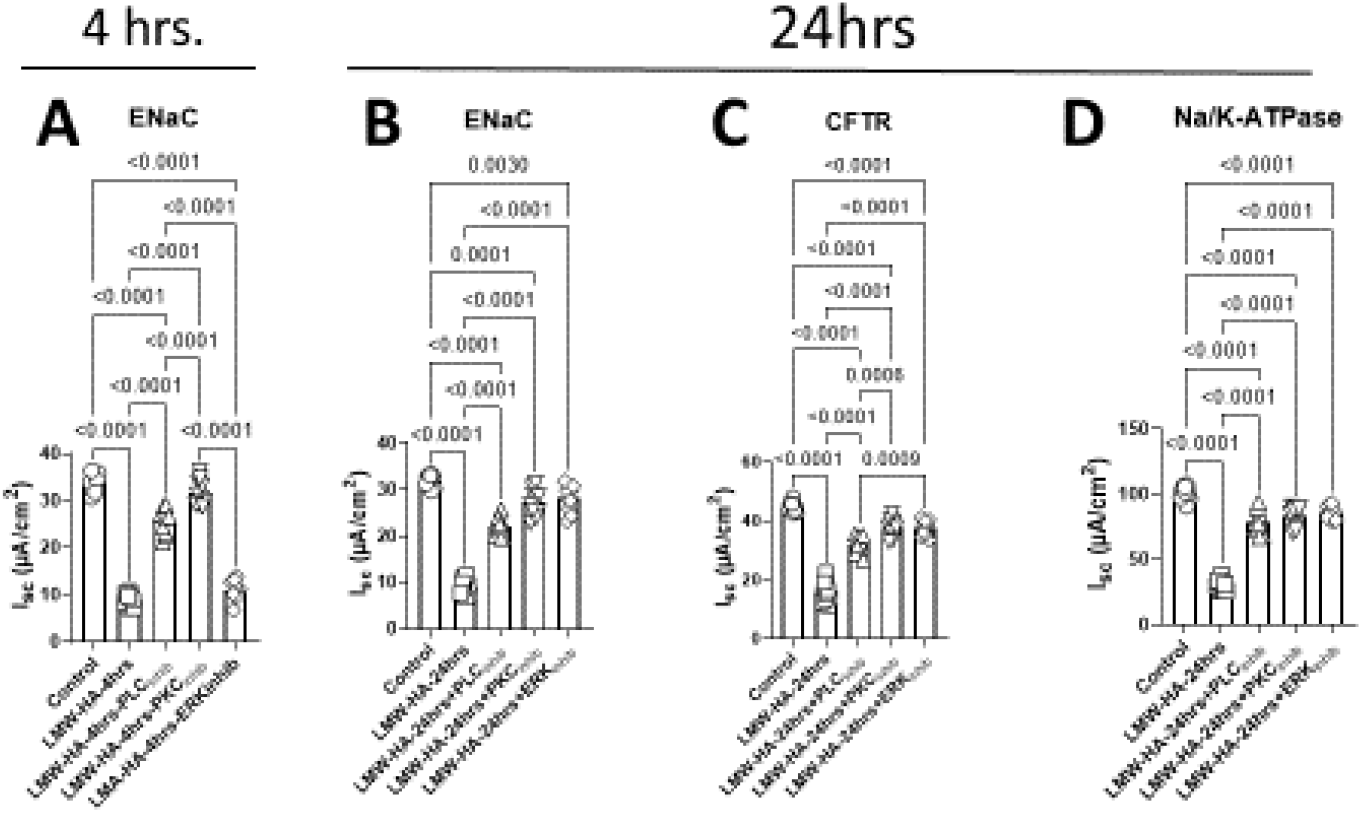
Application of PLC, PKC, and ERK inhibitors to MTEC monolayers 4hrs prior to cells exposure to 150ug/ml LMW-HA. **A**. 4hrs post LMW-HA application ENaC current was protected from LMW-HA when PLC or PKC were inhibited but not ERK. **B, C, and D**. 24hrs post LMA-HA apical application to MTEC monolayers, all currents were protected by inhibition of either PLC, PKC, or ERK. Data are Means ± SE, n=10. Significance was determined by 1-way ANOVA and post hoc Tukey test for multiple comparisons.

### CaSR is required for the activation of calcium-sensitive Cl^−^ channel by LMW-HA

To test the possibility that LMW-HA increases intracellular calcium via its action on the CaSR, we expressed CaSR in HEK-293 cells stably transfected with TMEM-16A, a calcium-sensitive chloride channel. **Figure 6F** shows the incubation of HEK cells with 150ug/ml LMW-HA activated Cl^−^ current through TMEM-16A in the presence CaSR (**Figure 6F**) but not when it was absent (**Figure 6B**). Furthermore, addition of 1μM NPS-2143 with 150μg/ml LMW-HA prevented the calcium-sensitive Cl^−^ current activation (**Figure 6G**). On the other hand, perfusion of both CaSR and mock transfected cells with Ringer’s containing 100μM ATP activated the calcium-sensitive Cl^−^ current through TMEM-16A **Figures 6D and 6H**. ATP acts on purinoreceptors to increase intracellular calcium and activate TMEM-16A independently of CaSR. The data suggest CaSR activation by LMW-HA is necessary for its action on lung ion channels.

**Figure 6.**
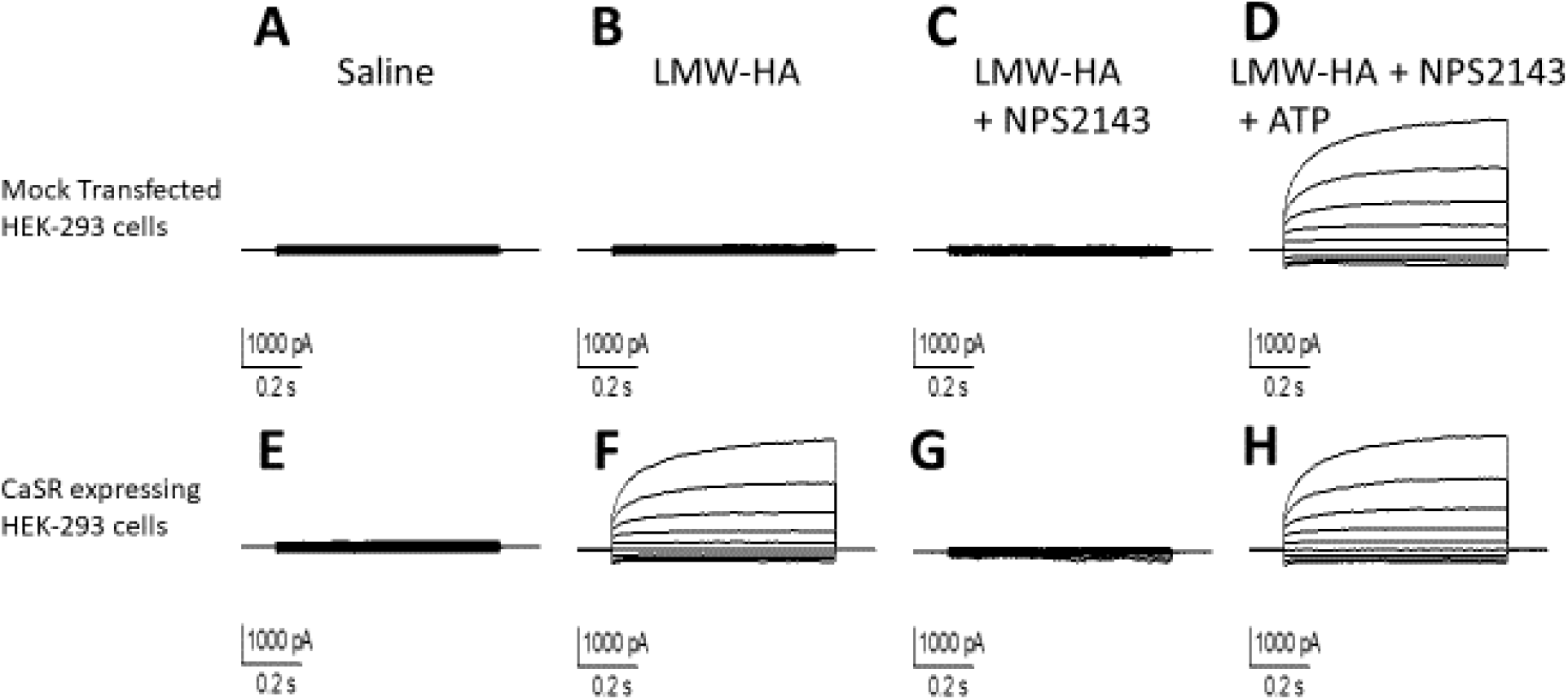
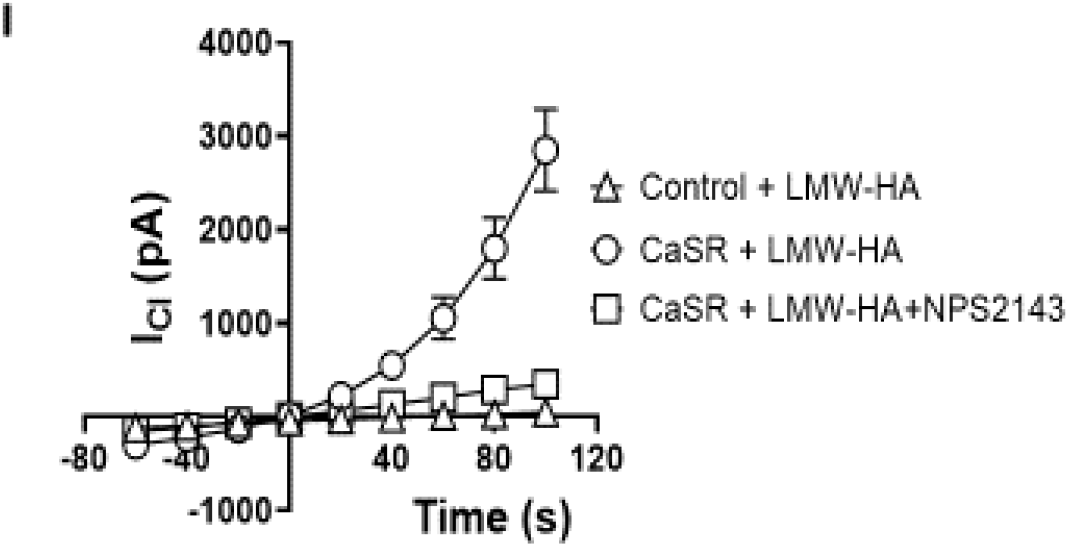
Whole cell patch clamp recording of Cl^−^ through TMEM-16 stably expressed in HEK-293 cells. **A**. Current recorded from a cell expressing TMEM-16 perfused with Ringer’s solution. **B**. Current recorded from a cell expressing TMEM-16 perfused with 150ug/ml LMW-HA in Ringer’s. **C**. Current recorded from a cell expressing TMEM-16 perfused with Ringer’s containing 150μg/ml LMW-HA and 1μM NPS-2143. **D**. Current recorded from a cell expressing TMEM-16 perfused with Ringer’s containing 150μg/ml LMW-HA, 1μM NPS2143, and 100uM ATP. ATP activated the calcium sensitive chloride current through TMEM-16 by increasing calcium in the cell. **E**. Current recorded from a cell co-expressing TMEM-16 and CaSR perfused with Ringer’s solution. **F**. Current recorded from a cell co-expressing TMEM-16 and CaSR perfused 150ug/ml LMW-HA in Ringer’s. LMW-HA activated the calcium sensitive Cl^−^ current through TMEM-16 by increasing calcium in the cell. **G**. Current recorded from a cell co-expressing TMEM-16 and CaSR perfused with Ringer’s solution containing 150μg/ml LMW-HA and 1μM NPS-2143. LMW-HA did not activate TMEM-16 because NPS-2143 inhibited CaSR. **H**. ATP activated the calcium sensitive Cl^−^ current through TMEM-16 by acting through its purinoreceptors. **I**. Current/voltage relationships of Cl^−^ current in HEK-293 cells expressing TMEM-16. LMW-HA activates the calcium sensitive Cl^−^ current only when cells are expressing both TMEM-16 and CaSR. The Cl^−^ current is inhibited when cells were perfused with 1μM NPS-2143 indicating the role of CaSR in intracellular calcium increase by LMW-HA.

### HMW-HA and NPS-2143 abrogate LMW-HA effect on mouse NPD

Previously we have shown that exposure of mice to chlorine (Cl_2_) increased the concentration of LMW-HA in their BAL and resulted in acute lung injury and airway hyperreactivity [19]. To investigate whether LMW-HA contributed to the inhibition of lung ion channels through its action on the CaSR in Cl_2_ toxicity, we exposed mice to Cl_2_ (400 ppm for 30 min) and 6hrs later we instilled them either with 50μl of 150μg/ml HMW-HA or 50μl of 1μM NP2143, and measured NPD at 24hrs post exposure to Cl_2_. As shown in, **Figure 7**, both agents reversed Cl_2_ induced inhibition on the amiloride sensitive Na^+^ driven NPD and the Forskolin activated GlyH101 inhibited Cl^−^ driven NPD. The role of CaSR in the injury inflicted by Cl_2_ inhalation is clearly evidenced by the reversal of NPD by more than 80% of the control values when 1μM NPS-2143 was instilled 6hrs post exposure to Cl_2_. Instillation of 50μl of 150μg/ml HMW-HA also reversed Cl_2_ induced injury, however, with unknown mechanism.

**Figure 7.**
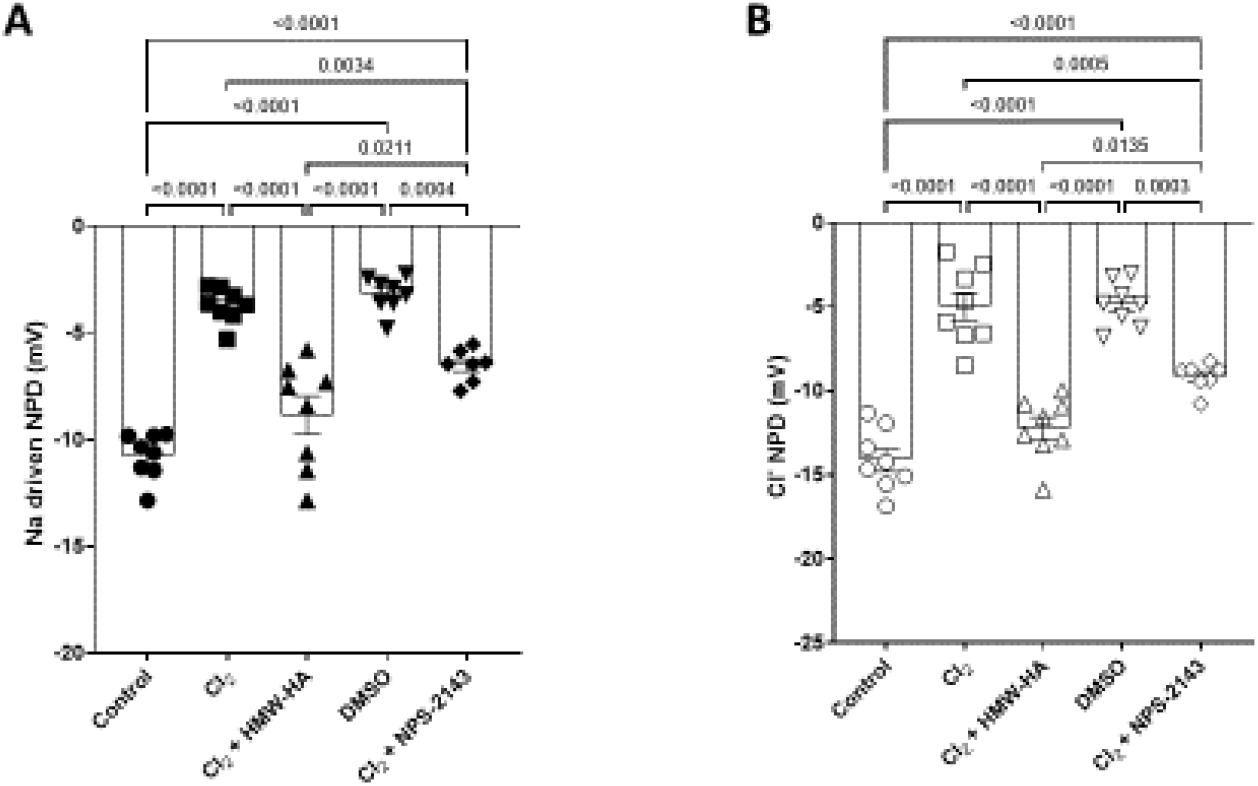
Nasal potential different was measured in control and chlorine exposed mice (400ppm for 30min). Cl_2_ inhalation reduced both the amiloride sensitive (**A**) and the Forskolin activated GlyH-101 inhibited (**B**) components by 64% and 69%, respectively, at 24hrs. Mice instillation with 50μl of 150μg/ml HMW-HA or 50μl of 1μM NPS-2143 at 6hrs post exposure reversed the effect of Cl_2_ on NPD to 90%, and 80% of control values, respectively. Data are means ± SE, n= 13-15, p<0.001. Significance was determined by 1-way ANOVA and post hoc Tukey test for multiple comparisons.

### LMW-HA inhibits ENaC activity in human ATII cells in-situ

In the next series of experiments, we incubated human lung slices (see methods section) with either saline of LMW-HA (150μg/ml) in Ringer’s solution for 4hrs and applied cell-attached mode of patch-clamp technique to measure ENaC activity in ATII cells, **Figure 8A**. Two distinct conductances are detected a 4pS and an 18pS. Slices incubation with 150μg/ml LMW-HA for 4hrs inhibited both ENaC conductances’, **Figure 8B**. 1μM NPS-2143 abrogated the effect of LMW-HA on 18pS conductance but not on 4pS conductance, **Figure 8C**. LMW-HA inhabited ENaC activity by decreasing both channels’ open probabilities (P_o_), 4pS, **Figure 8D**, and 18pS, **Figure 8E**, respectively. Inhibiting CaSR with 1μM NPS2143 restored 18pS activity and P_o_ by more than 80% of control P_o_, however, 4pS channel activity and P_o_ remained low. This is the first demonstration of ENaC function inhibition in human lung by LMW-HA activation of CaSR.

**Figure 8.**
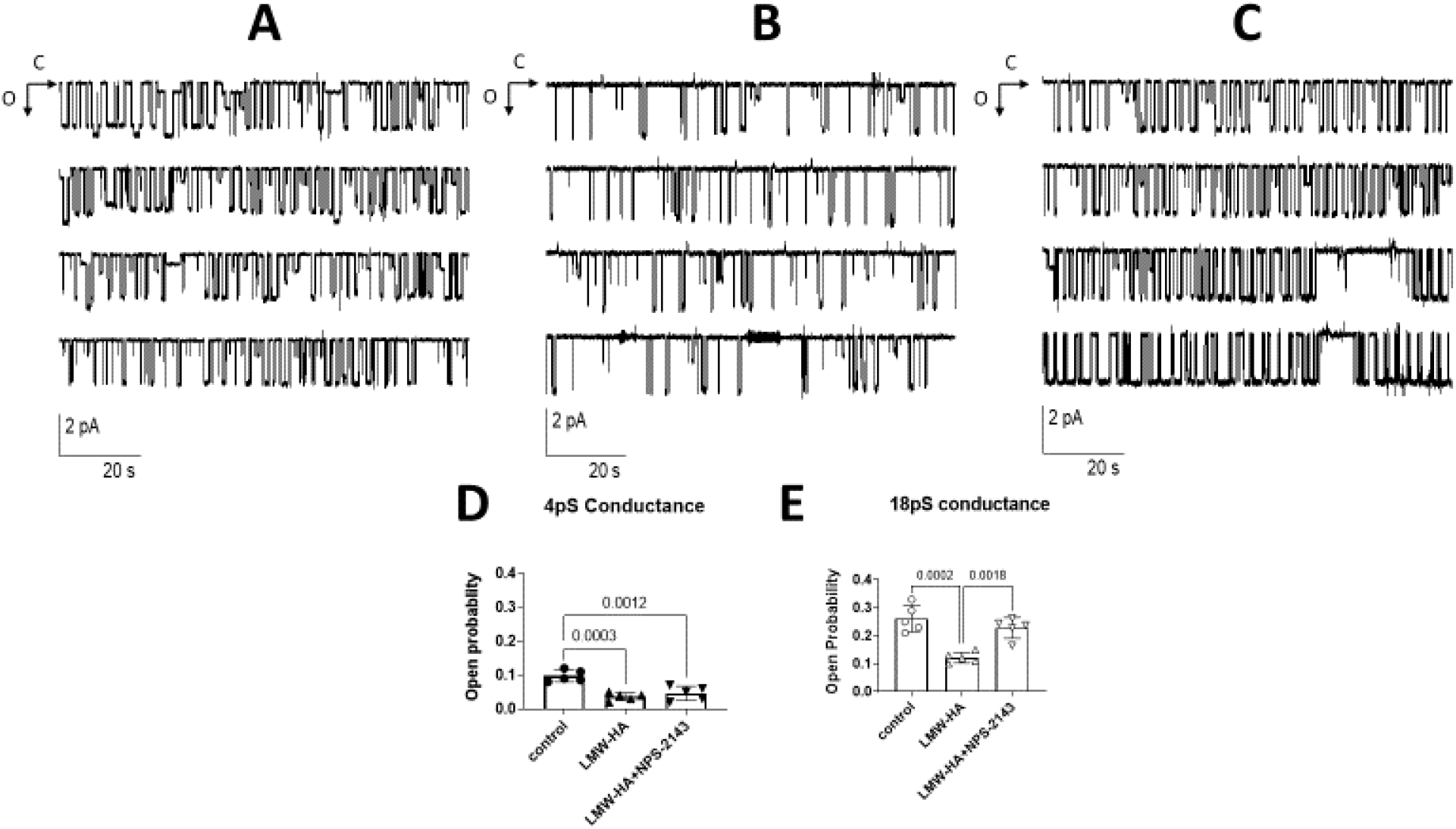
ENaC activity in human AT2 cells in-situ in precision cut lung slices from human lung tissue of a patient (more details are needed). **A**. Control recording from an ATII cells bathed in Ringer’s solution. Two conductance are visible, a 4pS and an 18 pS, the arrows indicate the open (o) and closed (c) states of the channels. **B**. Exposure of human lung slices to 150μg/ml LMW-HA for 4hrs reduced channels activities of conductances. **C**. incubation 1μM NPS-2143 abrogated LMW-HA effect on 18pS conductance but not on 4pS. **D** and **E** depict the summary data of both channels activities in control, 4hrs post exposure to 150μg/ml LMW-HA, and 4hrs post incubation with 150μg/ml LMW-HA and 1μM NPS-2143. The calcilytic prevented the effect of LMWHA on 18pS conductance but did not protect the 4pS conductance. Data are means ± SE, n= 13-15, p<0.001. Significance was determined by 1-way ANOVA and post hoc Tukey test for multiple comparisons.

## Discussion

In this study we demonstrate for the first time the modulation of epithelial ion and water transport across mouse and human respiratory epithelia by LMW-HA and CaSR. The activation of CaSR initiates an intracellular signaling cascade that stimulates PKC and inhibits ENaC function in human and mouse ATII cells, and AFC in mouse lung inducing lung edema at 4hrs post exposure to LMW-HA. The fact that LMW-HA can activate CaSR in epithelial cells and in airway smooth muscle cells and cause lung edema (this study) and AHR [19] provide new insight into the diverse effects of LMW-HA, the complex mechanisms of signaling through CaSR, and new concepts about lung edema formation and clearance following lung injury. In fact, nasal instillation of 1μM of NPS-2143 6hrs post mice exposure to 150μg/ml LMW-HA restored AFC and lung W/D ratio by reducing fluid accumulation in the lung. The inhibition of AFC in mouse lung as early as 4hrs revealed the speed at which the injurious LMW-HA fragments affect lung ion and water transport. Dynamic AFC process relies on the ability of lung epithelia to reabsorb Na^+^ ion through ENaC and drive the water with it to clear the air spaces of liquid. The inhibition of AFC at 4hrs coincides with the inhibition of ENaC function in human and mouse ATII cells in-situ and in MTEC monolayers at 4hrs post exposure to LMW-HA. Na^+^ absorption through ENaC is favored by the active transport of 3Na^+^ ion out of the cell by the Na,K-ATPase and the uptake of 2K^+^ ion. This process plays a major role in lung fluid clearance at birth and the maintenance of lung and alveolar spaces free of liquid thru life for effective gases exchange between inhaled air and blood. The effect of LMW-HA on ENaC as early as 4hrs without affecting the function of CFTR created a disequilibrium favoring liquid secretion and its accumulation in the lung as evidenced by lung W/D ratio, AFC inhibition, and the inability of the airways epithelium to reabsorb the accumulated liquid measured by μOCT.

Hyaluronan (HMW-HA) is an anionic, non-sulfated glycosaminoglycan polymer composed of D-glucuronic acid and N-acetyl-D-glucosamine, linked via alternating β-(1→4) and β-(1→3) glycosidic bonds. HMW-HA is uniquely characterized by its *de-novo* synthesis at the plasma membrane, it is extruded through the membrane into the extracellular matrix as it is synthesized by HA synthases (HAS1–3). Under physiological conditions, hyaluronan consists of 2,000–25,000 disaccharides, which corresponds to polymers of 2-25μm long and a molecular weight of 10^6^–10^7^ Dalton. HMW-HA interacts through hydrogen bonds, van der Waals, and electrostatic forces with specific proteins called hyaladherins and with membrane receptors, modulating development, morphogenesis, cell migration, apoptosis, cell survival, inflammation, and tumorigenesis [53]. It protects from ozone [32], and bleomycin injury [54, 55], smoke inhalation[56-58], sepsis [59, 60], and halogens inhalation toxicity[19, 61]. The breakdown of HMW-HA by hyaluronidases or reactive oxygen species generate smaller fragments of low molecular weight of less than 500 kDa that, by contrast to HMW-HA, promote inflammation, angiogenesis, epithelial to mesenchymal cell transition [62-66], and tissue injury [67, 68]. LMW-HA fragments stimulate cytokine production and activate the innate immune response by binding to CD44 that was found to be enhanced by inter-α-trypsin-inhibitor (IαI) [69, 70]. LMW-HA is also involved in airways hyperreactivity after exposure to ozone [32, 61], Cl_2_ and Br_2_ [19, 20, 61, 71], HCl [72], and various other forms of lung injury [67, 68, 73]. With so many functions attributed to hyaluronan, still little is known about its interaction with ion channels. In one study, hyaluronan was found to inhibit the pain sensitive channel, TRPV1, by direct interaction [16], and reduce the open probability of a stretch-activated ion channel heterologously expressed in oocytes[17]. It was found to induce lung hyper-reactivity by increasing Ca_i_^2+^ in airways smooth muscles through the activation of CaSR [20].

CaSR is predominantly expressed in the parathyroid glands and kidneys, where it regulates PTH secretion and renal tubular calcium re-absorption to maintain the external calcium concentration in the physiological range, 1.2 to 2.1mM, [21-23]. Initial studies of CaSR focused on its function in calciotropic tissues with obvious role in calcium homeostasis (parathyroid, kidney, and bone), but soon after CaSR was found expressed in different tissues with no apparent role in extracellular calcium homeostasis, including nervous system, lung, skin, placenta, breast, epithelia, and the endothelium [74]. Abnormal CaSR expression and function was found associated not only with calciotropic disorders, but also with diseases with-no obvious link to the calciotropic systems [25, 75, 76]. CaSR has been observed to control numerous cellular processes such as secretion, differentiation, proliferation, apoptosis, and gene expression including its own [24, 77].

The expression of the Ca-SR in the respiratory system plays an important role in fetal lung growth and development by stimulating fluid secretion in the pulmonary lumen [78, 79], lung branching, and morphogenesis in mice [80]. Ca-SR is expressed in human and mouse airway smooth muscle cells and bronchiolar epithelial cells [81], and its expression was increased in the vascular airway smooth muscle of patients with PAH [82, 83], asthma [25, 81, 84], allergen-sensitized mice [81], and in mice exposed to toxic chemicals such as chlorine and bromine [19, 20].

CaSR activation is initiated by the binding of a ligand and the activation of G-protein-dependent stimulation, via Gq/11, of phospholipase C (PLC) activity initiating the synthesis and the accumulation of IP_3_ and DAG. IP3 induces the release of Ca^2+^ from endoplasmic reticulum [85], while DGA activates PKC and MAP-kinases cascades and influence ion channels function and gene expression. Active PKC has been shown to downregulate ENaC function by phosphorylating specific amino acids on its subunits and predispose them to ubiquitination by Neural precursor cell-expressed developmentally down-regulated protein 4 (*Nedd4-2*) [86], initiating their internalization and degradation by the proteasome or lysosome. In this study we found the inhibition of CaSR with 1μM NPS-2143, to restore ENaC function in human and mouse ATII cells, AFC, ELF thickness measured by μOct, a finding that suggest the involvement of PKC in the inhibition of ENaC by CaSR activation 4hrs post exposure to LMW-HA.

Degradation of hyaluronan and its accumulation in the distal lung following injury was associated with edema formation [4, 5] because of its ability to attract and retain water. Our findings, however, suggest LMW-HA causes lung edema by inhibiting ENaC function through the activation of CaSR and its downstream kinases. ENaC function and fluid clearance were restored when LMW-HA was antagonized with HMW-HA or when CaSR function was inhibited with 1μM NPS-2143 at 6hrs post mice and cells exposure to LMW-HA. Similarly, the inhibition of PLC, PKC, or ERK prior to MTEC monolayers exposure to LMW-HA prevented ENaC inhibition at 4hrs and CFTR and Na,K-ATPase 24hrs.

I conclusion, the preservation of ELF thickness at ∼0.2μm suggests the involvement of exquisite homeostatic regulations by lung epithelia involving ENaC, CFTR, and Na,K-ATPase. Impairment of lung epithelia ion channels and Na,K-ATPase function leads to pulmonary edema [61, 87, 88], a serious complication triggered by many diseases and various types of traumata initiated by inhaled or aspirated particles, pathogens and toxic gases [89-92]. The precise occurrence and the associated morbidity and mortality of pulmonary edema are difficult to quantitate, as it is associated with such a wide variety of conditions. Thus, pulmonary edema is a major clinical problem for which there are no effective therapies to directly increase the rate of liquid absorption in the flooded lung. Cell matrix hyaluronan breakdown to smaller fragments of low molecular weight (<500 kDa) can cause lung injury or exacerbate preexisting conditions by activating intracellular signaling cascades downstream from CaSR, inhibiting epithelial ion and water transport leading to lung edema and ARDS. Antagonizing LMW-HA effects with HMW-HA or by inhibiting CaSR and the signaling cascades downstream from it using calcilytics may prove efficient in lung edema and ARDS treatment.

## Acknowledgments

Funding was provided by a REINVENT grant from the department of Anesthesiology and perioperative medicine to A. Lazrak and W. Song; CounterACT Program, National Institutes of Health Office of the Director, the National Institute of Neurological Disorders and Stroke, and the National Institute of Environmental Health Sciences, Grants 5UO1 ES026458, 3UO1 ES026458 03S1, and 1R21 ESo32956 to S. Matalon; NIH grant R21 AI152006 for A. Nellore and CW. Hoopes; NIH Grant R01 HL133006-0 to B. Woodworth, and P30 Grant to Gregory James Fleming Cystic Fibrosis Center of the University of Alabama at Birmingham.

